# BRAPH 2: a flexible, open-source, reproducible, community-oriented, easy-to-use framework for network analyses in neurosciences

**DOI:** 10.1101/2025.04.11.648455

**Authors:** Yu-Wei Chang, Blanca Zufiria-Gerbolés, Emiliano Gómez-Ruiz, Anna Canal-Garcia, Hang Zhao, Mite Mijalkov, Joana B. Pereira, Giovanni Volpe

**Author notes:** These authors contributed equally to this work (Yu-Wei Chang, Blanca Zufiria-Gerbolés).

## Abstract

As network analyses in neuroscience continue to grow in both complexity and size, flexible methods are urgently needed to provide unbiased, reproducible insights into brain function. BRAPH 2 is a versatile, open-source framework that meets this challenge by offering streamlined workflows for advanced statistical models and deep learning in a community-oriented environment. Through its Genesis compiler, users can build specialized distributions with custom pipelines, ensuring flexibility and scalability across diverse research domains. These powerful capabilities will ensure reproducibility and accelerate discoveries in neuroscience.

---

In recent years, network analyses have rapidly gained traction in neurosciences, linking multiple brain regions, clinical scores, and biomarkers to reveal system-wide interactions that often remain hidden when single variables are studied in isolation [1-4]. This approach is reshaping neurology, psychiatry, and cognitive sciences, driven by the promise of new tools for disease diagnosis, improved treatment monitoring, and deeper insights into how the brain functions. Here, we present BRAPH 2, a Flexible, Open-source, Reproducible, Community-oriented, and Easy-to-use (FORCE) framework for traceable and replicable network analyses. The standard BRAPH 2 distribution (https://github.com/braph-software/BRAPH-2) offers streamlined workflows for complex statistical analyses and deep learning, allowing researchers to study the dynamic interplay between brain structure, function, and behavior. Most importantly, the Genesis module of BRAPH 2 allows users to easily create and share custom distributions of BRAPH 2 with analysis pipelines fulfilling the FORCE principles, as illustrated in Figure 1. This unified approach will increase collaborations, accelerate innovation, and make advanced network analysis more accessible to researchers from neuroscience and related fields.

**Figure 1.**
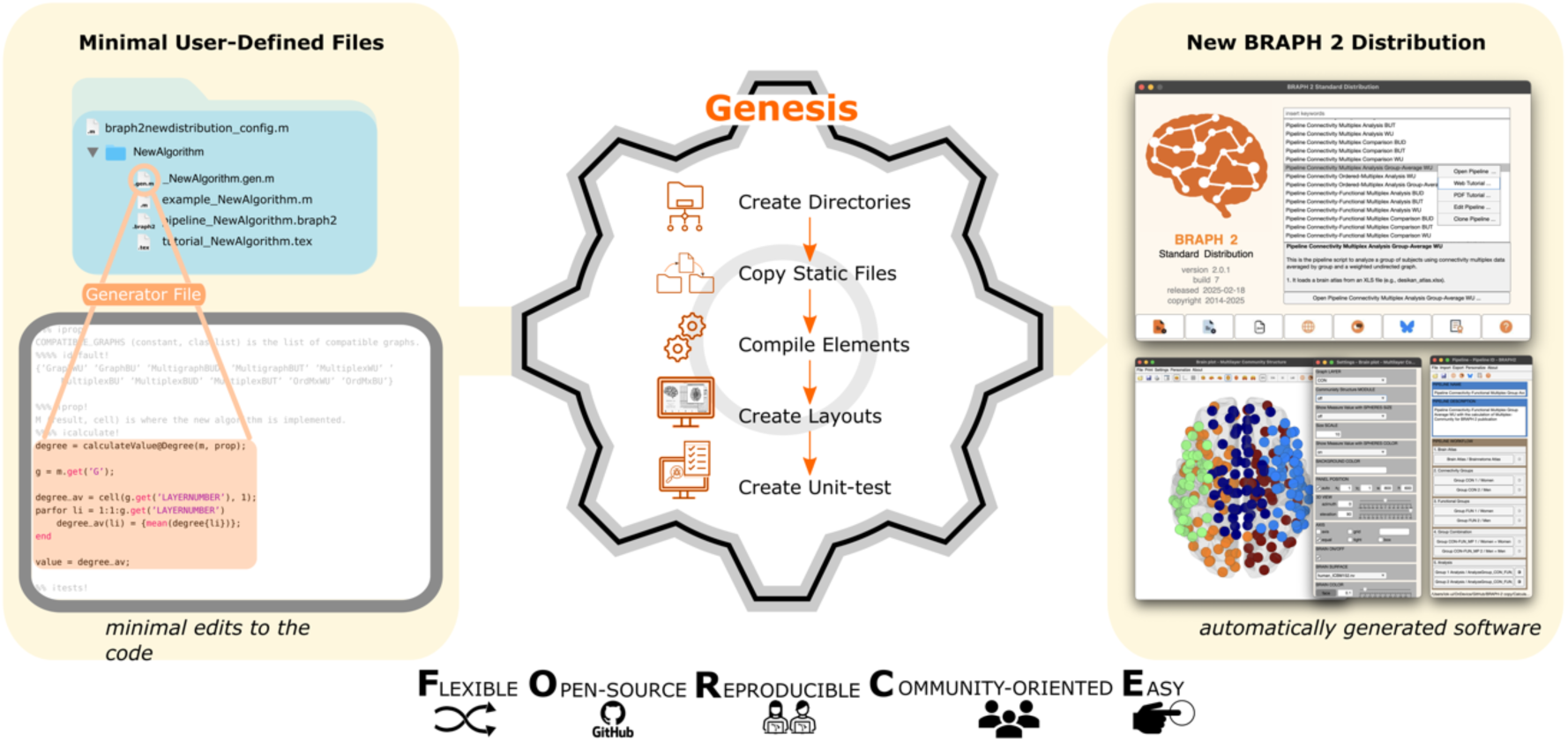
Genesis in action: compiling a custom BRAPH 2 distribution. Left: To create a specialized BRAPH 2 distribution with custom analysis pipelines, the user prepares a compilation configuration file (*braph2newdistribution_config*.*m*) and a folder with the necessary custom elements (e.g., *_NewAlgorithm*.*gen*.*m*), pipeline files (e.g., *pipeline_NewAlgorithm*.*braph2*), and optionally tutorial latex files (e.g., *tutorial_NewAlgorithm*.*tex*). Center: The Genesis module of BRAPH 2 compiles these files together with the core BRAPH 2 files in a series of steps including: creating directory structure, copying static files into the new directories, compiling elements, creating layouts for the graphical user interface (GUI), and generating unit-tests. Right: The custom BRAPH 2 distribution is compiled, complete with a no-code GUI for final users. This provides a framework for network analysis in neuroscience that is Flexible, Open-source, Reproducible, Community-oriented, and Easy-to-use.

The growing availability of large-scale, multimodal datasets has made network analysis increasingly complex [5-7]. Initiatives such as the UK Brain Biobank [8], the Human Connectome Project (HCP) [9], and the ENIGMA Consortium [10] have generated tens of thousands of high-resolution brain images, along with omics profiles and clinical records from diverse populations [5-7]. Although many software tools exist for handling these datasets, implementing new robust, reproducible, and traceable features remains a significant challenge [11]. These features require rigorous unit testing, the ability to trace outcomes back to original sources, continuous progress monitoring, and the development of user-friendly interfaces—tasks that often demand far more time and resources than creating core functionalities. Adopting standardized, containerized environments and automated testing frameworks can enhance reproducibility and traceability [12], but integrating these practices into research-oriented software is complex and time-consuming. As a result, research groups often prioritize the rapid development of new techniques at the expense of software quality, reproducibility, and long-term maintenance.

Many free and open-source software platforms for brain connectivity analysis exist, such as the Brain Connectivity Toolbox [13], eConnectome [14], GAT [15], CONN [16], BrainNet Viewer [17], GRETNA [18] and BRAPH 1 [19], among others. While these tools cover essential functions like importing data, computing network measures, and visualizing results, they often focus on specific tasks. This specialization often requires the use of multiple tools in various parts of the analysis, making it challenging to build comprehensive, full-stack workflows without human intervention prone to introducing mistakes and biases. Moreover, the rapid advancements in computational modelling and AI demand continuous software updates, which can delay the integration of the latest methods— especially in formats that are both reproducible and traceable. As a result, researchers face difficulties in adopting cutting-edge analyses in a manner that ensures consistency, transparency, and easy verification across studies, slowing scientific progress and potentially introducing mistakes.

BRAPH 2 addresses these challenges by introducing a user-friendly environment for building network analysis pipelines, complete with integrated statistical testing, reproducible and traceable research mechanisms, and a graphical user interface (GUI). By creating custom elements, users can expand both pipelines and GUIs, allowing the development of specialized software solutions without extensive coding. Moreover, being written in MATLAB, BRAPH 2 benefits from its cross-platform virtual environment that ensures portability across different operating systems. Though initially designed for brain imaging data, BRAPH 2 modular design supports a variety of data types. In fact, during its development, we created and tested specialized pipelines—such as those examining glycolytic synchronization waves in yeast cell populations [20] and exploring clustered dendritic spine loss in postmortem brain tissue [21]—not only as proof-of-concept cases but also to demonstrate BRAPH 2 flexibility for diverse research domains.

Compared to BRAPH 1 [19], BRAPH 2 has been completely rewritten for greater robustness and adaptability, addressing the need identified by users for more flexible but still reliable data analysis pipelines. At the heart of these improvements is the Genesis module of BRAPH 2, which uses simplified pseudocode to facilitate the creation and integration of new features and pipelines. The Genesis module permits the creation of custom BRAPH 2 distributions simplifying much of the underlying complexity related to developing and sharing robust software solutions. This will allow developers and users to adapt pipeline scripts and create new graphs, measures, or analyses with minimal programming effort. This streamlined approach ensures that custom elements integrate with the existing GUI and workflows, accelerating development while maintaining robustness. Detailed tutorials and developer guides are available, offering step-by-step instructions that illustrate how the Genesis module system supports a modular and expandable architecture. By simplifying the process of adding new methods and keeping the codebase maintainable, the Genesis module helps ensure that BRAPH 2 can rapidly incorporate cutting-edge techniques without sacrificing stability, reproducibility, or traceability.

The standard BRAPH 2 distribution exemplifies these ideas by including about 100 different pre-built analysis pipelines (https://github.com/braph-software/BRAPH-2). These pipelines include not only standard network analyses based on a single imaging modality but also multilayer graph analyses integrating diverse imaging and non-imaging datasets as well as deep learning pipelines. In Figure 2a, we demonstrate a use-case with pipelines available in the standard BRAPH 2 distribution, whereas Figures 2b and 2c showcase two other examples that highlight BRAPH 2 more advanced capabilities, which align with the criteria set forth by Neurodesk [12] or DeepPrep [22]. We describe these cases in detail below.

**Figure 2.**
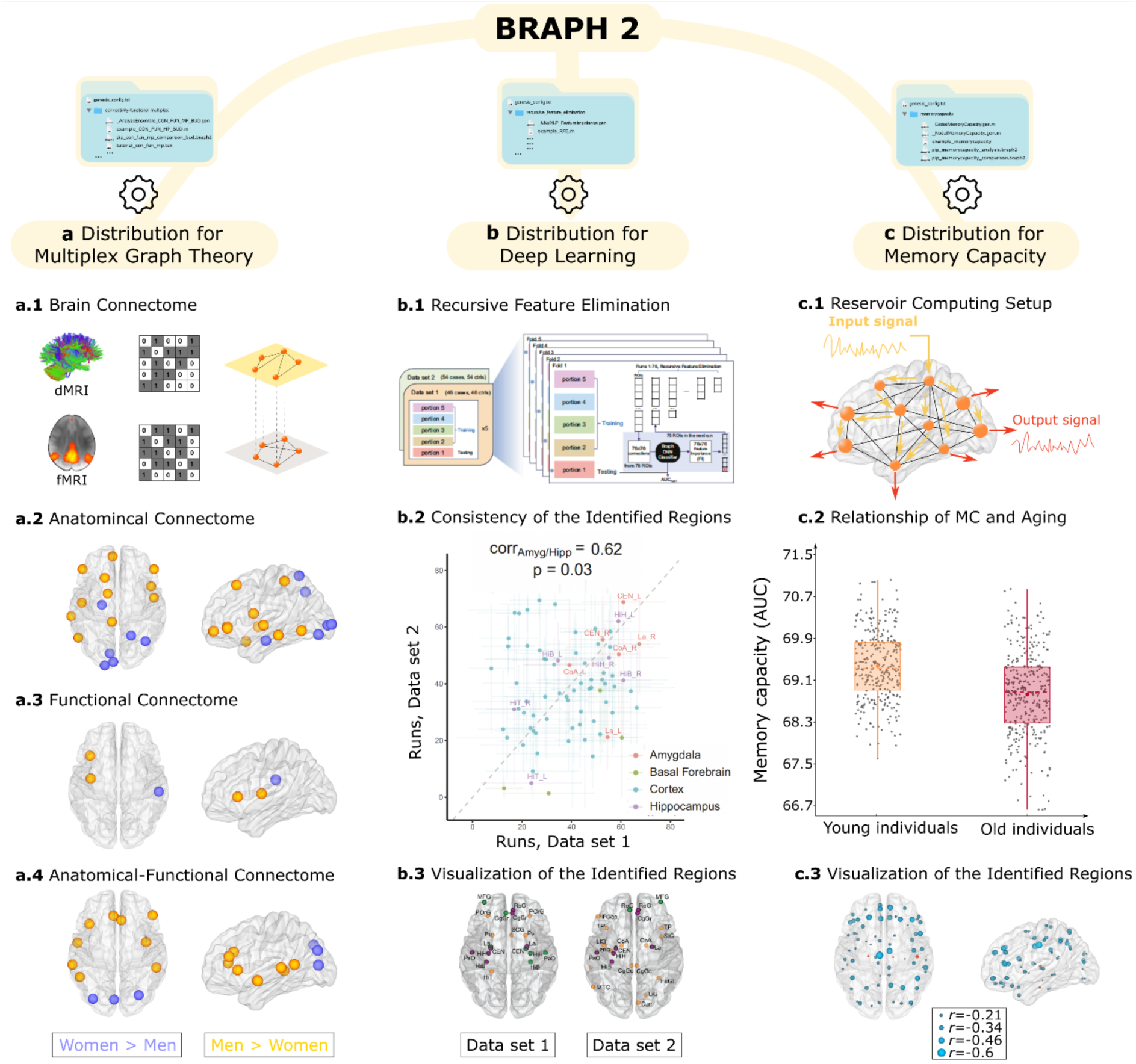
Examples of BRAPH 2 distributions. **a** An analysis pipeline from the Standard BRAPH 2 distribution. This pipeline is a multiplex graph theory, where anatomical and functional data are combined to create multiplex networks, providing deeper insights into brain organization. **b** Custom distribution used for deep learning focusing on recursive feature elimination, where a deep learning pipeline identifies consistent regions across two datasets using a multilayer perceptron and iterative feature elimination (used in [23]). **c** Distribution for memory capacity (MC) using reservoir computing, where memory capacity is estimated in an aging cohort; this demonstrates how a new algorithm can be provided to the community through a new BRAPH 2 distribution (used in [24], https://github.com/braph-software/MemoryCapacity).

In the first case study (Figure 2a), we used some analysis pipelines included in the standard BRAPH 2 distribution to analyze single-layer and two-layer multiplex networks derived from functional MRI (fMRI) and diffusion MRI (dMRI) data, focusing on finding sex differences in brain organization. A multiplex network is a network with several layers, where each layer represents one type of brain connectivity—in our case functional (fMRI) and anatomical (dMRI). By analyzing the two layers together, we gain insights into how functional and anatomical connectivity are related and interact with each other. Using this multiplex approach, we found that men and women show different core connectivity patterns—especially in the occipital, insular, frontal, and temporal brain regions— compared to single-layer (anatomical or functional only) graphs. The multiplex approach identified subtler differences that single-layer methods missed. This shows its potential for precision medicine approaches, where sex differences play an important role. Using a deep learning pipeline, we observed that the multiplex approach also improved classification accuracy, achieving an area under the curve (AUC) of about 0.80 for multilayer graph analyses compared to only 0.69 for single-layer approaches. By recording parameters and random seeds during statistical permutation tests, BRAPH 2 ensures that all randomization options and analysis steps are saved. This ensures reproducibility, allowing other users to rerun the exact same pipelines on different systems and obtain identical results. Moreover, BRAPH 2 supports data sharing by storing and sharing analysis pipelines without including sensitive personal data like age and sex, which are typically used only as covariates. This approach simplifies providing reproducible results in publications, which has become a standard requirement, and sharing data between research groups while maintaining privacy. Overall, this case demonstrates how users can achieve advanced analyses simply by running some BRAPH 2 ready-to-use pipelines provided with its standard distribution.

In the second case study (Figure 2b), we build on the functionalities of the standard BRAPH 2 distribution to investigate schizophrenia-related alterations in brain organization using two separate functional MRI datasets. These results are published in [23]. Here, we rely on the multilayer perceptron (MLP) classifier and feature-importance methods in the standard BRAPH 2 distribution, but we create an external recursive feature elimination script—iteratively training the MLP model, identifying the least important brain regions, removing them, and training the next MLP model—until we converge on a set of key regions linked to schizophrenia. Crucially, we find that the most important brain regions (e.g., hippocampus, amygdala, and specific cortical areas) overlap well across the two independent datasets and were also identified through an independent SNP-heritability enrichment analysis, providing evidence that they are robust disease-related hubs. By adapting a standard BRAPH 2 pipeline for classification with deep learning with a new iterative selection process, this example showcases how advanced users can expand built-in pipelines to suit novel research needs—still relying on the standard BRAPH 2 distribution while adding custom scripts.

The third case study (Figure 2c) shows the full power of the Genesis module by demonstrating how users can develop their own custom BRAPH 2 distributions. We demonstrate this for the quantification of the memory capacity of brain networks using reservoir computing across the lifespan [24]. Unlike the first two examples, where analyses build on standard BRAPH 2 pipelines, here we implement a custom algorithm and create a custom BRAPH 2 distribution with specialized pipelines. This distribution integrates the new memory capacity measure alongside all other needed features already available in BRAPH 2 (e.g., importing data, brain surface visualization, statistic modules). We apply this method to a lifespan cohort, showing that decline in computational memory capacity is a strong marker of aging. Moreover, our analysis finds that these measures correlate strongly with structural white-matter integrity, functional connectivity, and cognitive performance. By packaging the new pipeline into a standalone BRAPH 2 distribution, we ensure it can be easily shared with the community as a separate GitHub repository (https://github.com/braph-software/MemoryCapacity)—promoting reproducibility, portability, and rapid innovation.

Distributed as open-source software on GitHub, BRAPH 2 enforces high-quality standards through comprehensive unit tests, continuous integration, version control, and detailed documentation. Furthermore, it encourages researchers to modify and extend its source code, making it an ideal starting point for specialized analysis pipelines. Users can easily suggest additions, adapt existing workflows, and incorporate new modalities or analysis techniques to meet their specific research needs. The modular design and support for standard data formats simplify the process of adding new network measures, statistical tests, or visualization tools. This flexible foundation allows scientists to build on BRAPH 2 existing pipelines, rapidly expanding their functionalities while reusing and customizing proven methods. It runs on Windows, macOS, and Linux by using MATLAB’s virtual machine, as well as on hardware ranging from standard laptops to high-performance computing clusters. While memory requirements vary by analysis type, BRAPH 2’s efficient design scales well for large-scale data processing. Analyses and pipelines can be shared as MATLAB scripts or workflows. Furthermore, for those preferring a no-code experience, BRAPH 2 has a user-friendly GUI.

The Genesis module of BRAPH 2 allows developers to create and share state-of-the-art algorithms as custom BRAPH 2 distributions—without compromising the stability of the BRAPH 2 base distribution. New or experimental pipelines (e.g., the separate memory capacity repository: https://github.com/braph-software/MemoryCapacity) can be rapidly prototyped, tested, and refined.

This will strengthen BRAPH 2 as a dependable, flexible framework for researchers working across a wide range of scientific fields. Future development plans include integrating more advanced algorithms, multi-omics data tools, and additional deep learning methods (also taking advantage of the recent integration of MATLAB with Python [25], which will permit BRAPH 2 to use various Python deep learning frameworks natively) into the standard BRAPH 2 distribution. To maximize usability, BRAPH 2 provides extensive training resources and support, including user guides and tutorials available on its GitHub page (https://github.com/braph-software/BRAPH-2). This comprehensive online help and community forum will increase collaboration and knowledge sharing between both new and experienced users.

## Notes

https://github.com/braph-software/BRAPH-2

### Competing Interest Statement

The authors have declared no competing interest.

### Summary of Updates

- Correction of an author's name as previously misspelled. - Minor wording edits throughout the text to improve clarity and readability, without changing the scientific content.

